# Rapid and Programmable Protein Mutagenesis Using Plasmid Recombineering

**DOI:** 10.1101/124271

**Authors:** Sean A. Higgins, Sorel Ouonkap, David F. Savage

## Abstract

Comprehensive and programmable protein mutagenesis is critical for understanding structure-function relationships and improving protein function. However, current techniques enabling comprehensive protein mutagenesis are based on PCR and require *in vitro* reactions involving specialized protocols and reagents. This has complicated efforts to rapidly and reliably produce desired comprehensive protein libraries. Here we demonstrate that plasmid recombineering is a simple and robust *in vivo* method for the generation of protein mutants for both comprehensive library generation as well as programmable targeting of sequence space. Using the fluorescent protein iLOV as a model target, we build a complete mutagenesis library and find it to be specific and unbiased, detecting 99.8% of our intended mutations. We then develop a thermostability screen and utilize our comprehensive mutation data to rapidly construct a targeted and multiplexed library that identifies significantly improved variants, thus demonstrating rapid protein engineering in a simple one-pot protocol.

## INTRODUCTION

Directed mutagenesis of a desired protein is an important technique both for understanding structure-function relationships as well as improving protein function for research, biotechnology, and medical applications. For example, techniques like deep mutational scanning, where every position in a protein is mutated to all possible amino acids, can be applied to understand key variants associated with disease^1^, while targeted mutagenesis of proteins such as Green Fluorescent Protein (GFP) have expanded our capacity to visualize many biological processes^2^. The ability to generate comprehensive mutation libraries and programmed libraries focused on specific locations or amino acids is crucial to these applications.

In order to address these needs, many polymerase chain reaction (PCR) based approaches have been developed. Firnberg and Ostermeier have built libraries composed almost entirely of single mutations using specialized protocols based on uracil-containing template DNA^3^, while Melnikov and Mikkelsen constructed a comprehensive library by splitting one gene into many different regions small enough to be synthesized on a programmable microarray, followed by multiplexed *in vitro* recombination^4^. Belsare and Lewis have demonstrated targeted, combinatorial library construction using alternating cycles of fragment and joining PCR^5^. Nevertheless, all these techniques suffer from the requirement of multiple complex *in vitro* reactions that are labor-intensive and require specialized protocols or reagents.

An alternative approach would be to incorporate synthetic oligonucleotides *in vivo* directly into a gene of interest in a programmable fashion. In *E. coli*, oligonucleotides introduced into the cell via electroporation can recombine with the genome or resident plasmids with the help of the lambda phage protein Beta, in a process termed recombineering^6^. Mechanistically, it is thought that Beta-bound oligonucleotides anneal to the replication fork of replicating deoxyribonucleic acid (DNA) and are subsequently incorporated into the daughter strand, thus directly encoding mutations into a new DNA molecule^7^. Recombineering is therefore a compelling method for genetic manipulation. Cheap and easily obtained standard oligonucleotides are the only varying input and the protocol - mixing oligonucleotides in one-pot reactions - is straightforward.

This process was shown to be capable of mutating the *E. coli* genome for rapid metabolic engineering in a process termed Multiplexed Automated Genome Engineering (MAGE), which used multiple rounds of recombineering to increase the penetrance of mutations^8^. Other work has demonstrated that thousands of pooled, barcoded oligonucleotides can be used, in parallel, to modify the expression of > 95% of *E. coli* genes and map their effect on fitness^9^. More recent studies have combined recombineering with the programmable DNA nuclease Cas9, as a means of enforcing mutational penetrance, to mutate tens of thousands of loci in parallel with high efficiency^10^.

Despite its success in genome engineering, recombineering of plasmids is relatively uncharacterized^11^. Plasmid recombineering (PR) is of particular interest in protein mutagenesis as plasmids are easily shuttled between different strains and organisms for cloning and screening. Notably, recombineering strains achieve enhanced mutation efficiency by knocking out mismatch repair and possess a higher genome-wide mutation rate^12^, which can complicate screens or selections sensitive to suppressor mutations. The use of plasmids, however, uncouples protein variation from any background mutation in the genome. Thomason et al. have previously demonstrated that PR is capable of generating mutations, insertions, and deletions with efficiencies comparable to genomic recombineering. We reasoned that the principles of MAGE – a one-pot reaction and multiple mutation rounds – would be applicable to PR as well.

To benchmark comprehensive PR for protein engineering we sought to measure the efficiency, bias, and overall performance of saturation mutagenesis on the small protein iLOV. iLOV is a 110 residue protein derived from the Light, Oxygen, Voltage (LOV) domains of the *A. thaliana* phototropin 2 protein^13^. The native LOV domain binds flavin mononucleotide (FMN) and uses this co-factor as a photosensor to direct downstream signal transduction. Mutational analysis has revealed that a cysteine to alanine substitution in the FMN binding site interrupts the native photocycle and instead dramatically increases the protein’s fluorescent properties. iLOV is an ideal candidate for further engineering because fluorescent proteins that don’t require molecular oxygen for chromophore maturation are a desirable alternative to green fluorescent protein. In previous experiments, DNA shuffling was used to isolate iLOV, a variant that has six amino acid mutations relative to the wild-type (WT) phototropin 2 LOV2 sequence and an improved fluorescence quantum yield of 0.44^13^. Additional approaches to engineer further improved iLOV variants have also relied on error-prone PCR and DNA shuffling, missing much of the possible sequence-space^14^. Due to the potential utility of iLOV and its comparatively limited engineering relative to other fluorescent proteins, we hypothesized iLOV could serve as an excellent model system for exploring the utility of PR. Finally, the gene length of iLOV is exceptionally short (330 bp) and analysis of iLOV libraries is suited to deep sequencing. Current paired-end sequencing covers the entirety of the open reading frame and can accurately identify all mutations to a single sequenced plasmid. This provides insight into the mechanisms and utility of recombineering.

Here we demonstrate that PR is capable of constructing both comprehensive protein libraries and targeted mutagenesis libraries focusing on a small section of sequence space. We built a complete mutagenesis library of iLOV and found it to be specific and unbiased, detecting 99.8 % of our intended mutations. We explored this fitness landscape in the context of thermostability using a plated-based screen that allowed us to identify many desirable thermostabilizing mutations. To demonstrate the iterative and programmable nature of our platform, we designed and built a multiplexed library focused on these mutational hotspots and isolated significantly more stable variants. In total, this work demonstrates that plasmid recombineering is a rapid and robust method for the generation of protein mutants for both unbiased, comprehensive libraries and programmable targeting of specific regions in sequence space.

## RESULTS AND DISCUSSION

### A one-day, one-pot reaction generates a comprehensive mutation library

To generate a comprehensive mutation library of iLOV, oligonucleotides were designed to target the gene within a high copy ColE1 plasmid (Supp. Figure 1). The target plasmid contained a promoterless iLOV coding region to prevent growth biases between mutants possessing different fitness. 10 ng of target plasmid was mixed with an equimolar mixture of 110 recombineering oligonucleotides, one primer for each codon in iLOV (Figure 1A). These oligonucleotides were 60 bp long and complementary to the lagging strand, which was previously demonstrated to be more efficient than targeting the leading strand^15^. Oligos contained a centrally located NNM mutation codon, (where N = A/C/G/T and M = A/C) which encodes all amino acids except methionine and tryptophan. Tryptophan, in particular, is known to quench flavin fluorescence in flavoproteins^16^ and was excluded from the library. Although modified oligonucleotides have been shown to enhance recombineering efficiency, e.g. phosphorothioation, standard oligonucleotides were used to minimize cost and complexity^8^.

**Figure 1.**
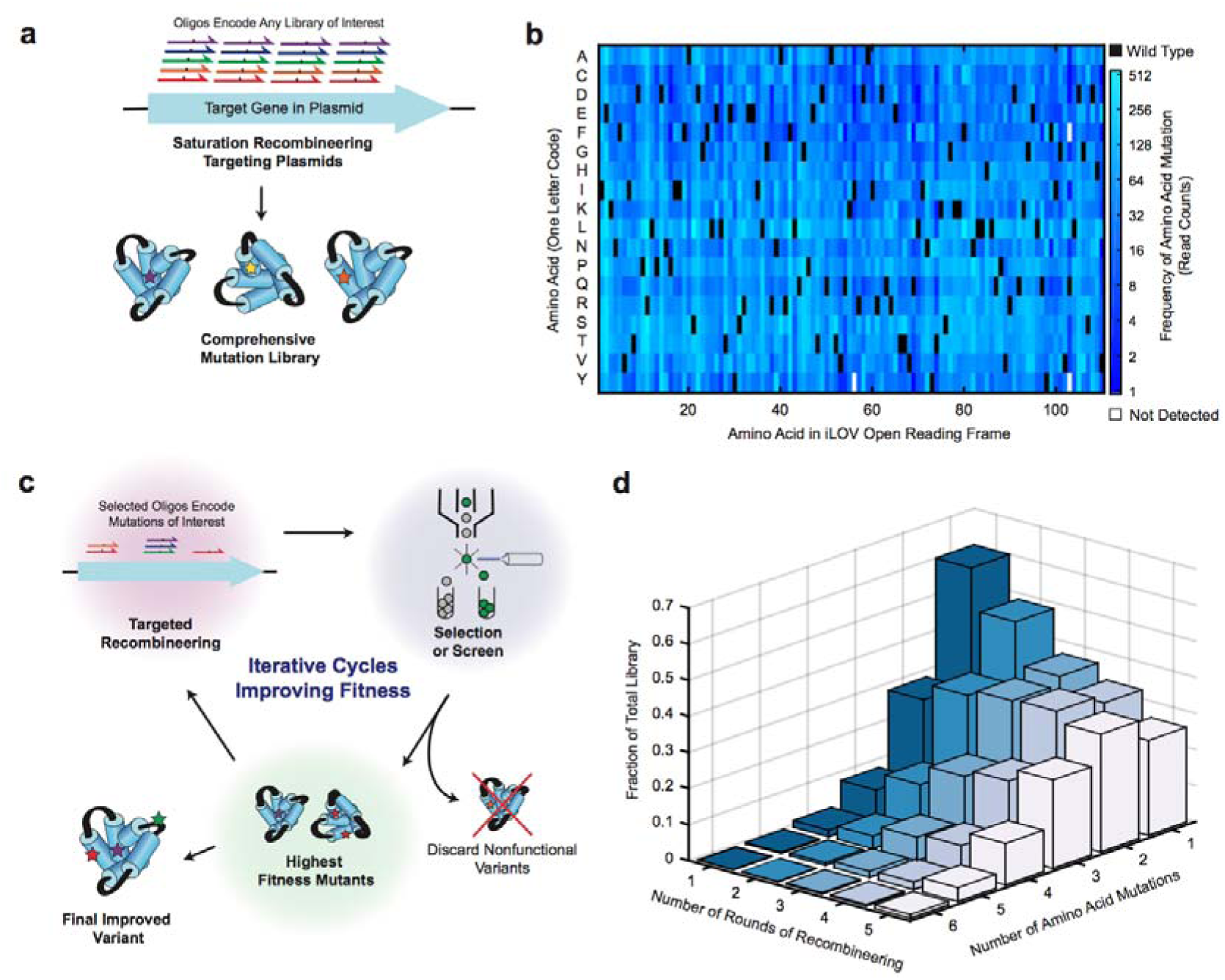
Plasmid Recombineering (PR) of iLOV generates a specific and comprehensive mutation library. (A) Cartoon of comprehensive PR accomplished using synthetic oligonucleotides tiled across the target gene. (B) Frequency of single amino acid mutations mapped by residue and location in the Round 1 library. Black indicates the WT iLOV residue. White indicates no detected reads. (C) Cartoon of programmable PR targeting a specific sequence space. Sampling of the highest fitness mutants can inform subsequent library design, with recombineering oligos specifically targeting mutations of interest. (D) Mutational distribution of the library after additional rounds of recombineering.

Initial experiments confirmed that PR can be used to generate diverse libraries in a programmable fashion. The plasmid and oligonucleotide mixture was first electroporated into the recombineering strain EcNR2^8^, grown overnight and miniprepped. Deep sequencing of the recovered library, hereafter termed the Round 1 library, revealed substantial mutagenesis compared to the non-recombineered plasmid (Supp. Figure 2). Further analysis revealed that, while the majority of reads were WT iLOV sequence, 29% of reads contained a single codon mutation. These mutations covered every position in the protein and nearly all targeted amino acid conversions were observed (Figure 1B). 1867 out of the possible 1870 single residue mutations were detected in the Round 1 library (169972 reads passing quality threshold).

Further founds of PR were used to increase the penetrance of mutations. Although the Round 1 library covered targeted mutations comprehensively, it was roughly 61% WT iLOV sequence. Furthermore, the programmable nature of PR allows the construction of more targeted libraries with combinatorial multiplexing of high fitness mutations, such as would be useful in directed evolution (Figure 1C). Four additional rounds of recombineering were performed to reduce the WT fraction of the library and investigate the distribution of variants with 2+ mutations. The number of reads with codon mutations increased substantially with further rounds of recombineering (Figure 1D). In the Round 5 library, single mutations were the most common (33%), with the remainder composed of WT sequences (26%) and sequences containing 2+ mutations (41%).

### Recombineering libraries can be finely controlled to alter the composition of mutations

In the absence of prior information, the ideal mutagenesis technique would produce every mutant targeted with equal frequency. Because each variant is initially present in the same amount, a uniform library requires the least amount of screening (or selection) in order to isolate improved mutants. A non-uniform method, in contrast, might produce a highly variable distribution of mutants and requires more screening or selection in order to fully explore sequence space. We therefore analyzed our sequencing data to characterize the uniformity and sequence preferences of PR libraries.

We detected two distinct types of bias: positional bias and mismatch bias. In positional bias, oligonucleotides targeted to different positions in the coding sequence incorporate with characteristically different efficiencies (regardless of the mutation produced at that position). In mismatch bias, oligonucleotides that are more similar to the WT sequence (e.g. differing only at the first base of the targeted codon) incorporated with high efficiency while more divergent oligos (e.g. different at all three positions) incorporated with lower efficiency. Positional bias is evident when comparing the frequencies of single codon mutations across all 110 codons in iLOV (Figure 2A). Despite the nearly 60 bp of homology between the oligonucleotide and the template plasmid, oligos targeting adjacent codons can exhibit a 2-3 fold different incorporation. The largest such discrepancy is found at codon 42, which is mutated 5.7 times more frequently than codon 41. Variation in efficiency has been observed in other recombineering studies and was somewhat correlated with the oligonucleotide binding energy^8^. While our data was not clearly correlated with binding energy (Supp. Figure 3), a biological replicate revealed replicable positional bias (Supp. Figure 4), suggesting the presence of an underlying physical mechanism for positional bias.

**Figure 2.**
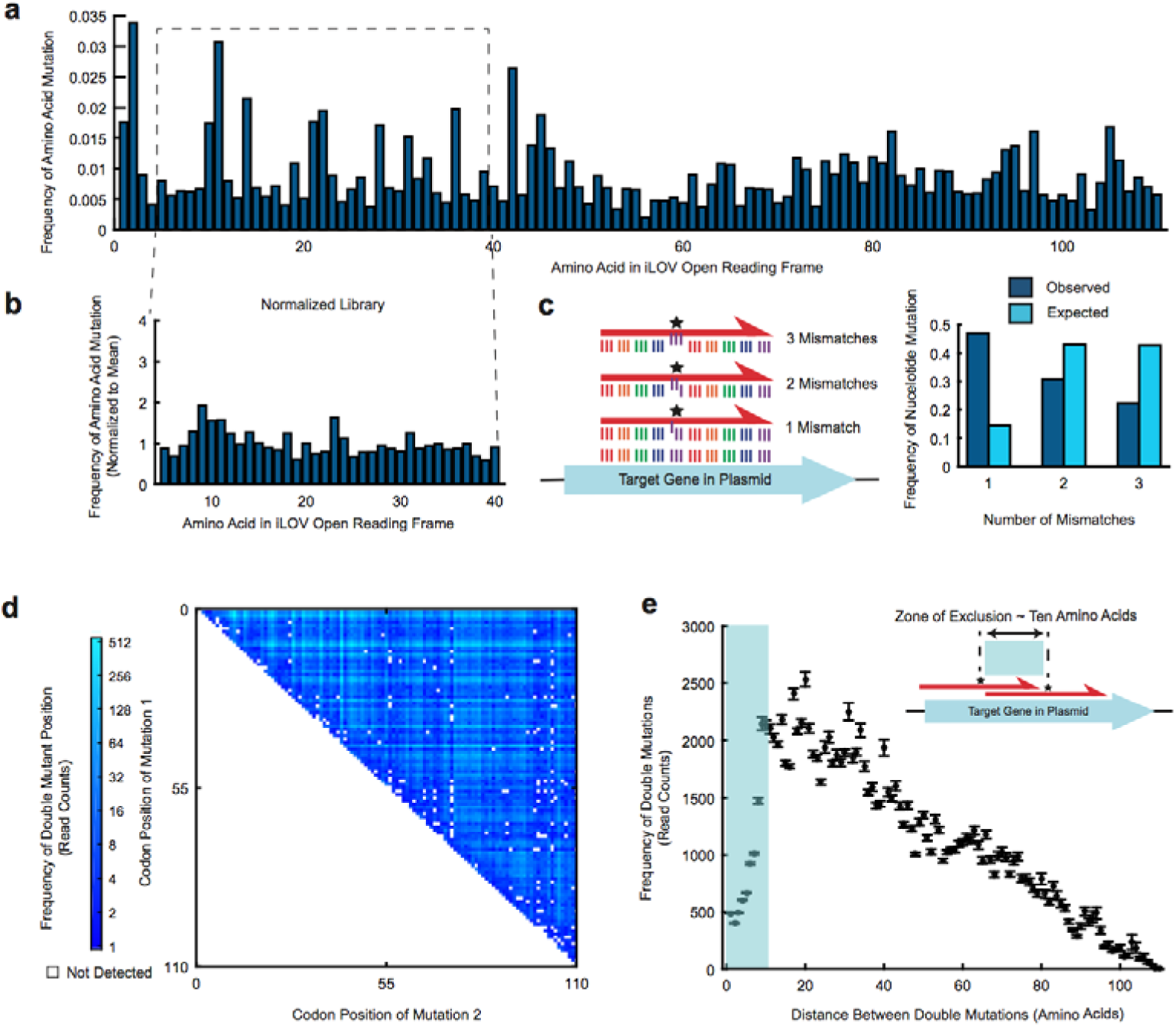
The recombineering mechanism produces moderate bias within the library and can be manipulated by varying the delivered oligonucleotide concentration. (A) Frequency of single amino acid mutations across iLOV in the Round 1 library. (B) Biological replicate of Round 1 library with normalized oligonucleotide concentrations. (C) Observed vs. expected distribution of one, two, or three nucleotide mismatches among single amino acid mutations. The 95% confidence intervals for these measurements are all smaller than +/− 0.0025 (normal approximation to the binomial distribution). (D) Frequency of double mutations in the Round 5 library. White = not detected. Highly represented rows and columns arise from positional bias (Supp. Figure 6). (E) Absolute read counts in the Round 5 library as a function of the pairwise distance between double mutations. Note the decreasing linear trend arises from fewer possibilities of double mutations with increasing distance. Error bars, standard deviation.

Comprehensive mutagenesis applications, such as deep mutational scanning, ideally begin with uniformly distributed mutants in a naïve library. We hypothesized that the positional bias observed in the Round 1 library – constructed using equimolar mutagenic oligonucleotides – could be corrected by altering the mixture of oligonucleotides used in the electroporation step. A second library was therefore constructed by normalizing the concentration of each oligonucleotide in the library according to the frequency of mutations obtained at the corresponding position in Round 1. This library was constructed using PR, and the first 40 amino acids were deep sequenced. The normalized library was indeed more uniform, with a largest adjacent codon discrepancy of 2.1 fold, compared to 3.3 fold in the equimolar replication library (Figure 2B).

In order to understand the importance of this normalization in a quantitative fashion, we performed a simulation to evaluate the pragmatic impact of oligonucleotide normalization on effective library size – the number of samples that must be taken from a library to achieve a desired representation of the library diversity. In the ideal case, i.e. mutants are found in the library with equal frequency, if one wishes to sample, say, 95 % of the diversity contained within the library, we must screen roughly three times the number of distinct members. That is, the effective library size is threefold the targeted library size. Our equimolar and normalized libraries are not uniformly distributed, but random sampling from our sequenced data *in silico* allows calculation of the effective library size. Our simulation revealed a substantial improvement in sampling efficiency, from a mean effective library size of 139 in the original library to 107 in the normalized library, which is quite close to that of an ideal uniformly distributed library (Supp. Figure 5).

The presence of mismatch bias in the library demonstrates that recombineering favors incorporation of oligos that are more similar to the WT template sequence. Our oligonucleotides all contained one, two, or three nucleotide mismatches relative to WT iLOV. After computationally enumerating all possible oligonucleotides for each codon in iLOV, we expected that 14% of oligonucleotides would contain one mismatch, 43% two, and 43% three. In contrast, oligonucleotide-template pairs with two or three mismatches were observed at only 31% and 22% respectively, while single nucleotide mismatches were overrepresented by more than threefold at 47% (Figure 2C).

While a single round of PR generates many single mutants, additional rounds shift the distribution to increasing mutation numbers. The Round 5 library, for example, is 25% double mutants. These mutations are well represented, with 5817 out of a possible 5940 locations detected (Figure 2D). However, it is clear that the same positional bias seen in the Round 1 library is preserved in subsequent rounds. This is to be expected if recombineering events are statistically independent, and can be seen by the correlation of double-mutation hotspots with single mutation positional bias (Supp. Figure 6). This data suggests that double mutations in a normalized library will be far more uniformly distributed.

The double mutation data also indicates that some recombineering events are not perfectly independent. A plot of the pairwise distance between all detected double mutants reveals an uneven distribution in frequency (Figure 2E). Specifically, double mutations are less likely to be within 30 bp (i.e. 10 amino acids) of one another. This effect becomes more pronounced with additional rounds of recombineering (Supp. Figure 7). In the Round 5 library, this ‘zone of exclusion’ is significant enough that double mutations a few amino acids apart are nearly four times less frequent than double mutations with a much larger separation (e.g. three amino acid gap versus 10 amino acid gap). We hypothesize that this bias is due to the mechanism of recombineering which requires oligonucleotides to anneal to a complementary locus via homology arms flanking the NNM mutagenic codon. In this mechanism, sequential oligonucleotides would ‘overwrite’ previous mutations due to incorporation of the most recent oligonucleotide’s homology arms, which extend 30 bp on either side of the central NNM. Another potential mechanism that disfavors incorporation of nearby double mutants is mutational reduction of oligo incorporation efficiency. In this hypothetical mechanism, mutations generated in early rounds of PR could reduce the homology and, therefore, incorporation efficiency of oligos in subsequent PR rounds.

### Screening the iLOV library identifies mutations conferring thermostability

Previous screening and structural work indicates that a well-packed binding site for the FMN fluorophore may lead to improved photochemical properties of iLOV, such as photostability, by limiting the dynamics of the FMN chromophore and its ability to dissipate energy following excitation^14^. Additionally, searches for improved LOV-based fluorescent reporters have turned up homologous variants such as CreiLOV that, while brighter (50% greater quantum yield), exhibit substantial toxicity upon expression^17^. More generally, Tawfik and colleagues have theorized that thermostable proteins serve as more fruitful starting points for engineering and directed evolution^18^. In this view, variants with greater thermostability can better tolerate mutations, increasing the likelihood of observing mutations that improve protein function in a manner unrelated to thermostability. We thus investigated the thermostability of our library, which contained nearly every single amino acid mutation and a small fraction of possible double mutations.

We created a plate-based assay to screen for thermostabilized variants in the Round 5 library. The library was plated at high colony density onto standard LB-Agar, grown overnight, then incubated at 60 °C for two hours (Figure 3A). This treatment completely abrogates the fluorescence of WT iLOV as well as nearly all mutants (Figure 3B). Some colonies remained fluorescent, however, and these were recovered and expressed in 96-well plate format. Cultures expressing these library members were lysed, clarified, and analyzed for thermostability. Fluorescence measurements were taken every 0.5° C during a temperature ramp from 25° C to 95° C and used to calculate the melting temperature (T_m_) for each clone, the temperature at which 50% of maximal fluorescence is retained (Figure 3C). All 93 assayed clones demonstrated a substantial increase in thermostability relative to iLOV (Figure 3D). Improvements in T_m_ ranged from two to nearly ten degrees C.

**Figure 3.**
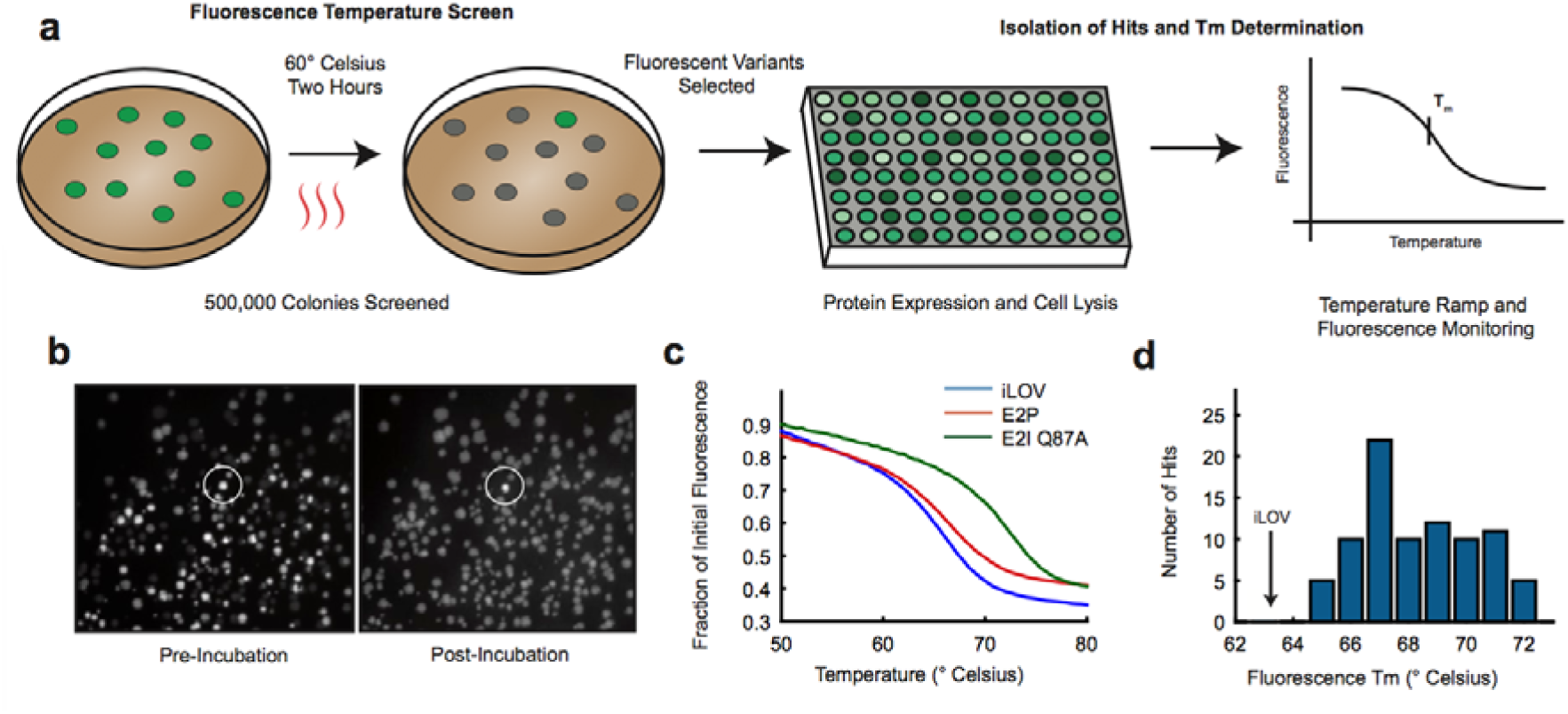
A plate-based thermostability screen identifies mutations that improve iLOV fluorescence at elevated temperatures. (A) Cartoon of the thermostability screen assay and subsequent hit validation procedure. (B) Representative fluorescent images of library colonies before and after temperature challenge. (C) Representative fluorescent thermal melt curves of lysate for two thermostable hits. (D) Histogram of T_m_s for 93 thermostabilized iLOV variants.

### Recombineering based multiplexing allows rapid and robust directed evolution

Many methods now exist for generating comprehensive single mutant libraries. However, most such methods are incapable of building targeted libraries for exploring the effect of mutations at sites of interest highlighted by previous mutational or structural studies. Because PR is capable of both targeted and comprehensive mutagenesis, we hypothesized that PR could fulfill these requirements in a cost-effective and straightforward one-pot, one-transformation protocol.

To demonstrate the rapid multiplexing capability of PR, we designed a second library containing the top 25 most frequent thermostabilizing mutations from the initial plate screen (Figure 4A). Several of these mutations consisted of alternative amino acids at the same position, and in these cases a different oligonucleotide was designed for each. The encoding oligonucleotides were designed such that homology arms would stop short of neighboring mutations so as not to overwrite them (Supp. Figure 8).

**Figure 4.**
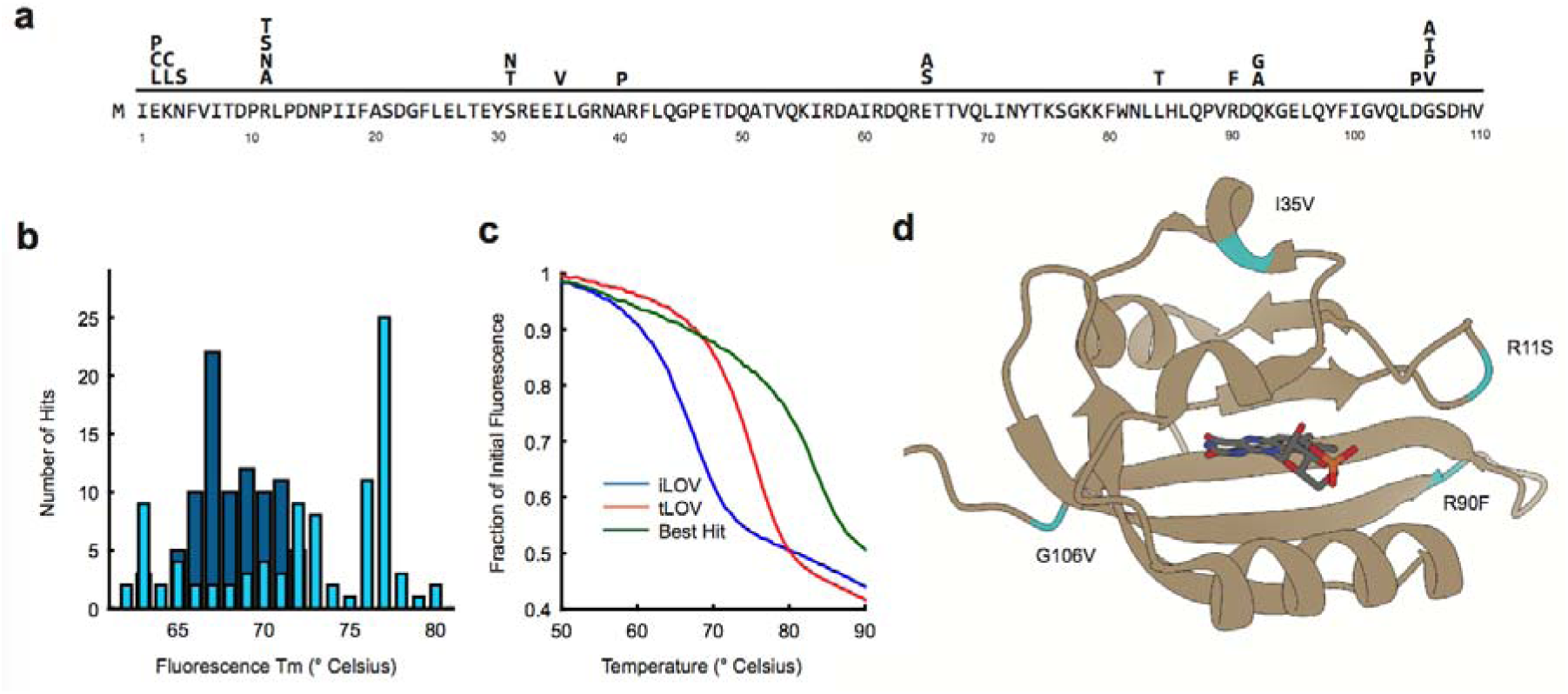
Multiplexing thermostabilizing mutations rapidly identifies doubly improved iLOV variants. (A) 25 thermostabilizing mutations mapped to the iLOV protein sequence. (B) Histogram of initial thermostable hits (Blue) superimposed on hits from the multiplexed library (Cyan). (C) Representative melt curves of two thermostabilized variants compared to iLOV. (D) Crystal structure of iLOV (PDB 4EES) indicating the location of four mutations found in tLOV: R11S, I35V, R90F, G106V.

This multiplexed library resulted in striking improvements to thermostability, with the best variants having T_m_ values nearly 20° C greater than iLOV (Figure 4B). Isolating and re-cloning these variants verified that despite the significant increase in T_m_s, the shape of their melt curves was not significantly different from that of iLOV, even for the most thermostable mutant (Figure 4C). It was noted during the screening process that one variant in particular, hereafter referred to as thermostable LOV (tLOV), seemed to produce abnormally bright lysate under high expression conditions. tLOV and iLOV were expressed and purified in parallel and their absorbance and emission were characterized *in vitro*. To accurately quantify relative quantum yield, the emission curves were normalized for absorption at 450 nm and integrated. tLOV was found to be approximately 10% brighter (Supp. Figure 9), and sequencing revealed the presence of four mutations scattered throughout the protein, all introduced by PR (Figure 4D). Notably, none of these locations are directly within the FMN binding pocket, and their contribution to thermostability or quantum yield improvement are not obvious. Thus, it would have been difficult to rationally design tLOV. Moreover, a targeted mutagenesis technique is required for isolating quadruple mutants reliably: iLOV is a small protein, but a quadruple mutant library would contain > 10^13^ variants, well beyond our current screening capacities. Finally, thermostability was verified by differential scanning calorimetry of iLOV vs tLOV (Supp. Figure 10).

An ideal method for generating comprehensive protein libraries would be simple and robust, enabling both complete and targeted mutagenesis without a change in reagents. In this study, we demonstrate that PR can be effectively used for both comprehensive and programmable mutagenesis of the fluorescent protein iLOV, using methods that are generalizable to any gene of interest.

Previous recombineering studies have successfully built libraries in the genome, demonstrating specificity and fine control of mutational composition. Wang et al.^8^ used multiple rounds of recombineering to mutagenize six consecutive nucleotides using 90 bp oligonucleotides and found a 75% mutation rate after five rounds. We find quite similar behavior in the sequencing analysis of the iLOV recombineered plasmid libraries constructed here. Our Round 5 library consisted of 74% mutant variants, and the mutations were well distributed in sequence space. The modest positional bias could be substantially ameliorated by altering the ratios of oligonucleotides added to the electroporation mixture. This resulted in a nearly uniform distribution of mutations such that the effective library size was almost ideal (Supp. Figure 5). This approach could easily be used to accommodate more complex libraries with weighted mutation frequencies at various locations. The correlation of replicate libraries (Supp. Figure 4) strongly suggests an underlying physical mechanism for differing oligonucleotide recombineering efficiencies, and it will be important to understand this effect in order to predict efficiencies rather than rely on empirical data as used here. Regardless of these modifications, the reagent cost and experimental effort remain minimal in all cases – standard 60 bp oligonucleotides, a one-pot reaction, and simple cycles of electroporation, growth, and plasmid isolation.

Additional biases resulting from the mechanism of recombineering were detected and while their magnitude was smaller, their effect on library size and screening can be significant. Previous work has found that recombineering efficiency drops sharply with the size of the modification made^8^. Here, too, single nucleotide mutations were observed to be more common than double and triple nucleotide mutations. This effect alters the distribution of codons present in the final library, with the template codons determining this frequency shift. Notably, mismatch bias is intrinsic to the annealing of oligonucleotides and is likely present for *in vitro* methods as well.

The ‘zone of exclusion’ around a first mutation generates another mode of bias. A second mutation is less likely to appear inside this zone than outside of it. This effect became more pronounced with increasing rounds of recombineering. Is it likely this results from oligonucleotides’ homology arms overwriting earlier mutations during later rounds of PR. In other words, we hypothesized that homology arms are capable of introducing revertant mutations. We thus predict that the length of homology arms would impact the length of the ‘zone of exclusion.’ As some amount of homology is absolutely required for recombineering, no form of recombineering is suitable for efficiently generating adjacent mutations from different oligonucleotides. This limitation can be overcome by using single oligonucleotides encoding sequential mutations but at the cost of reduced efficiency due to mismatch bias. Again, if this effect fundamentally stems from the annealing of oligonucleotides then it is likely present for *in vitro* methods as well.

*In vitro* methods represent the most powerful alternatives for generating comprehensive mutation libraries. The chief advantage of PCR-based mutagenesis is that it can generate a library composed almost entirely of single mutations. Ostermeier and colleagues accomplished this by utilizing specialized protocols based on uracil-containing template DNA^3^. Such methods have been used to generate nearly comprehensive libraries of genes for exploring the entirety of the fitness landscape^19^. The final proportion of wild-type sequences in this work is comparable to that of the Round 5 library (∼25%). In contrast, the remainder of their library is composed almost entirely of single mutations only, while our Round 5 library contained 41% 2+ mutation variants. These two approaches thus generate very different mutation profiles, and choice of one or another will depend on the goals of any particular experiment.

In directed protein evolution, iterative rounds of mutagenesis can be used to multiplex fitness-improving mutations. PCR-based protocols^3^, direct gene synthesis^4^, and some other recombineering techniques^10^ excel at generating libraries composed of single mutations. However, in many applications, 2+ mutations are desired at non-contiguous locations. Recent work has developed a PCR-based method to accomplish this goal *in vitro*^5^, and we hypothesized that PR was well suited to serve as a complementary approach *in vivo*, doing away with cloning altogether. To this end, we comprehensively explored the iLOV single mutation sequence-space for thermostability, selected the fitness enhancing mutations, and demonstrated the utility of PR for advanced protein engineering by multiplexing many different single and double mutations at discontinuous sites across iLOV in a second library. The ability to select and easily mutate numerous specific and non-contiguous locations across a protein is highly useful for a variety of techniques that utilize experimental or phylogenetic data to computationally predict and enhance enzymes^20^, explore epistatic interactions^21^, or even scan SNPs in human proteins for disease prediction^1^.

iLOV engineering has been relatively limited in comparison to other fluorescent proteins^13^. Because the domain has been taken out of its natural structural context, we hypothesized that its thermal stability could be increased. Consistent with this idea, many mutations were found to improve the thermal stability of iLOV up to a robust, 10° C increase in T_m_. These improvements were then stacked by multiplexing the 25 most common mutations from the first screen. As a measure of convenience, the same one-pot, single electroporation protocol was used, and the library was designed, built, and tested in one week. One particular variant among the thermostabilized pool, tLOV, was found to be ∼10% brighter that iLOV *in vitro*. This result is consistent with previous work demonstrating that constraining the FMN fluorophore can improve the photochemical properties of iLOV^14^. It would be interesting to perform comprehensive mutagenesis of the thermostabilized iLOV mutants in search of further improvements to the protein’s brightness or red/blue spectral shifting, as increased thermostability has been hypothesized to permit greater exploration of function-altering mutations^18^.

In summary, we have demonstrated that PR retains many of the ideal properties of genome recombineering, including specificity and programmability. We found that PR was suited for the construction of both comprehensive and targeted libraries, and that the simplicity of the protocol led to rapid and reliable screening experiments. In particular, PR is suitable for cycles of iterative design, construction, and sampling of genetic libraries, requiring no specialized reagents or protocols. We developed a thermostability screen of the fluorescent protein iLOV and used the resulting mutation data to rapidly construct a multiplexed library that identified significantly improved variants, including the first enhancement to the protein’s brightness since its development.

## MATERIALS AND METHODS

### Strains and Media

Strain EcNR2 (Addgene ID: 26931)^8^ was used for generating PR libraries in plasmid pSAH031 (Addgene ID: 90330). For thermostability screening and protein expression, iLOV libraries were cloned into pTKEI-Dest (Addgene ID: 79784)^22^ using Golden Gate cloning^23^ with restriction enzyme BsmbI (NEB) and transformed into either Tuner (Novagen) or XJ b Autolysis *E. coli* (Zymo Research). Unless otherwise stated, strains were grown in standard LB (Teknova) supplemented with kanamycin (Fisher) at 60 μg/mL.

### Recombineering Library Construction

Libraries were constructed using a modified protocol from Wang 2011^24^. Briefly, 110 oligonucleotides (Supp. Table 1) or 25 thermostabilizing oligonucleotides (Supp. Table 2) were mixed and diluted in water. A final volume of 50 μL of 2 μM oligonucleotides, plus 10 ng of pSAH031, was electroporated into 1 mL of induced and washed EcNR2 using a 1 mm electroporation cuvette (BioRad GenePulser). A Harvard Apparatus ECM 630 Electroporation System was used with settings 1800 kV, 200 Ω, 25 μF. Three replicate electroporations were performed, then individually allowed to recover at 30° C for 2 hr in 1 mL of SOC (Teknova) without antibiotic. LB and kanamycin was then added to 6 mL final volume and grown overnight. Cultures were miniprepped (QIAprep Spin Miniprep Kit) and monomer plasmids were isolated by agarose gel electrophoresis and gel extraction (QIAquick Gel Extraction Kit) to remove multimer plasmids^11^. The three replicates were then combined, completing a round of PR.

### Library Sequencing and Analysis

The iLOV open reading frame was amplified from PR libraries by PCR to add indices and priming sequences for deep sequencing (Supp. Table 3). PCR products were sequenced on Illumina platform sequencers (MiSeq and HiSeq) through the Berkeley Genomics Sequencing Laboratory. Sequencing data were analyzed with a custom MATLAB pipeline. Briefly, reads were filtered to remove those that were of low quality, frameshifted, or did not exactly match the annealing portion of the amplifying primers. An internal control (Supp. Figure 2) verified the low error rate of PCR and Illumina sequencing relative to PR. Finally, full-length reads were compared with the iLOV target sequence for mutation analysis.

### *In silico* Effective Library Size Simulation

Simulations were performed using a custom MATLAB script. The simulations focused on amino acid positions 5 – 40, for which we possess comprehensive frequency data from the equimolar oligonucleotide and normalized PR experiments. Successive sampling, with replacement, of these 36 positions was performed, with each position’s likelihood commensurate with the observed mutation frequency. Once 34 out of 36 positions had been observed in the simulation, which we arbitrarily define as ‘well-sampled’ (94.4% of the library diversity), the total number of samples taken was recorded as the effective library size. This process was then repeated 10^4^ times for each library to generate a distribution of effective library sizes (Supp. Figure 5).

### Thermostability Screening

Colonies were screened on 10 cm dishes containing standard LB-agar with kanamycin. Tuner cells expressing the library were found to be fluorescent in the absence of induction after 24 hours. Plates were incubated at 60° C for 2 hours, after which the vast majority of colonies were no longer fluorescent. Approximately 500,000 colonies were screened, of which 244 remained fluorescent. These colonies were pooled, miniprepped, and deep sequenced to identify protein mutations. This DNA was also transformed into XJb cells to recover individual variants. Colonies were then grown overnight in 96-well deep well plates with 1 mL of LB + kanamycin supplemented with 100 μM Isopropyl β-D-1-thiogalactopyranoside (IPTG) and 3 mM arabinose at 37° C. Cultures were frozen and thawed to lyse the cells, then clarified by centrifugation. Supernatant was analyzed in a StepOnePlus^™^ Real-Time PCR System (Applied Biosystems) to estimate a T_m_ for each variant.

### Protein Expression and *in vitro* Characterization

iLOV variants were expressed and lysed in XJb cells as above but in 100 mL volume. Excess FMN (Sigma) was added to the lysate to ensure all proteins possessed ligand. iLOV variants were then purified by nickel affinity chromatography using HisPur^™^ Ni-NTA Resin (Thermo Scientific). Variants were filtered using Vivaspin 6 3,000 molecular weight cut-off (Sartorius) with phosphate buffered saline (PBS) pH 7.4 (Gibco). Variants were further purified by size exclusion chromatography using a NGC Chromatography System (Bio-Rad). Purified proteins were stored at 4° C in PBS. For quantum yield determination, emission between 460 nm and 600 nm was measured in a FluoroLog Spectrophotometer (Horiba), and 450 nm absorbance was measured in an Infinity M1000 PRO monochromator (Tecan). Emission curves were integrated in MATLAB and normalized for absorbance. Thermostability measurements were made in a Nano differential scanning calorimeter (TA Instruments).

## SUPPORTING INFORMATION

The Supporting Information is available free of charge on the ACS Publications website at DOI:

Figure S1: Map of iLOV PR target plasmid. Figure S2: Internal control for sequence analysis pipeline. Figure S3: Recombineering efficiency correlation with binding energy or hairpin formation. Figure S4: Recombineering efficiency correlation between replicates. Figure S5: Simulation of effective library size *in silico*. Figure S6: Positional bias correlation with double mutations. Figure S7: ‘Zone of Exclusion’ for Rounds 1,3, and 5. Figure S8: Design of multiplexed library oligonucleotides. Figure S9: Emission scan of purified iLOV and tLOV. Figure S10: Thermostability of iLOV and tLOV via differential scanning calorimetry. Table S1: PR oligonucleotides for comprehensive iLOV library. Table S2: PR oligonucleotides for multiplexed thermostability library. Table S3: PCR primers used for Illumina sequencing. Table S4: Nucleotide and amino acid sequences of iLOV and tLOV.

## AUTHOR INFORMATION

### Corresponding Author

. Address: 2151 Berkeley Way, Berkeley, CA 94704; (510) 643-7847

### Author Contributions

S.A.H. and D.F.S designed the research. S.A.H. and S.O. performed the experiments. S.A.H. performed the computational analysis and analyzed the data. S.A.H. and D.F.S wrote the paper. Reagents described in this work are available on Addgene https://www.addgene.org/David_Savage/).

### Notes

The authors declare no competing financial interest.

## ACKNOWLEDGMENTS

We thank H. Wang (Columbia) for providing the *E. coli* strain EcNR2. We would like to thank A. Flamholz and B. Oakes for productive discussions and readings of the manuscript, and C. Cassidy-Amstutz for assistance with *in vitro* biochemistry. This work was supported by NIH New Innovator award 1DP2EB018658-01 to D.F.S., and S.A.H. was supported by NIH Training Grant 5T32GM066698-10 and Agilent.

